# A simple, reversible and non-toxic anchor-away system for effective nuclear depletion of proteins

**DOI:** 10.1101/2025.11.28.691147

**Authors:** Sofia Esteban-Serna, Tove Widén, Hanne Grosemans, Iseabail Farquhar, Sander Granneman

**Affiliations:** Centre for Engineering Biology, University of Edinburgh, Edinburgh, UK; Department of Life Sciences, Chalmers University of Technology, Gothenburg, Sweden; Centre for Cell Biology, University of Edinburgh, Edinburgh, UK; School of Biological Sciences, University of Edinburgh, Edinburgh, UK

**Keywords:** Conditional depletion systems, anchor-away, chemically-induced dimerisation, abscisic acid, rapamycin

## Abstract

The anchor-away (AA) technique enables rapid depletion of nuclear proteins by tethering them to cytoplasmic anchors through rapamycin-induced heterodimerisation. Albeit powerful, in *Saccharomyces cerevisiae,* this system is restricted to rapamycin-resistant strains, as the drug inhibits TOR signalling and hinders the heat-shock response, limiting its application to stress-related studies. Moreover, this AA method is not fully reversible, limiting studies of dynamic cellular processes that require transient perturbation and functional restoration. To overcome these constraints, we developed an alternative AA system that uses the plant hormone abscisic acid (ABA) to induce conditional association of the target to its cytoplasmic anchor. The ABA-AA system enables efficient and fully reversible depletion of highly abundant nuclear proteins. Unlike rapamycin, ABA does not cause major gene expression changes and is suitable for diverse genetic backgrounds. The ABA-AA system provides a fully reversible, non-toxic, and broadly applicable alternative for nuclear protein depletion across eukaryotic systems.

## Introduction

Common approaches to characterising protein function in yeast include permanently knocking out the gene encoding it or generating a conditional allele (e.g., a point mutation that yields a temperature-sensitive protein, which is functional at a permissive temperature but loses function at a non-permissive, warmer temperature). However, essential genes cannot be deleted or inactivated, and generating temperature-sensitive mutants can be labour-intensive, often leading to slow loss of function of the protein of interest. Moreover, in the latter case, the required temperature increases can trigger heat stress responses, which can overshadow the adaptive response caused by the deactivation of the protein of interest.

A popular alternative strategy to conditional gene knockouts is genetic depletion, which can be mediated by promoters or riboswitches. In most instances, endogenous promoters are replaced with inducible ones. Typical conditional promoters include the native *pGAL* and *pMET*, which are turned on by glucose and methionine, respectively ^1,2^; or heterologous promoters that are repressed in the presence of doxycycline/tetracycline ^3–5^. Similarly, riboswitches inserted into the 5′ untranslated region (UTR) of the target gene can, upon ligand binding, block ribosome scanning and thus modulate translation initiation of the gene product in a controlled manner ^6–8^. Disadvantages of these conditional depletion systems is that they often involve altering culture nutrient conditions and long incubation periods to achieve sufficient depletion of the target protein.

In contrast, engineered chemically induced dimerisation (CID) methodologies can often achieve significant and rapid depletion of the target protein, often in less than 30 minutes, thereby minimising noise introduced by secondary responses. CID systems employ inducers, which are small membrane-permeable molecules that promote stable interactions between two distinct protein domains. Such is the mode of action underlying the two most broadly used CID variants, namely the auxin-inducible degron (AID) and the anchor-away (AA) methods ^9–15^ (Fig. 1A). In the AID system, the target is tagged with an AID epitope that binds the rice protein TIR1 in the presence of auxin (Fig. 1A). In turn, TIR1 recruits ubiquitin ligases to polyubiquitinate the protein of interest, marking it for proteasomal degradation (Fig. 1A).

**Figure 1.**
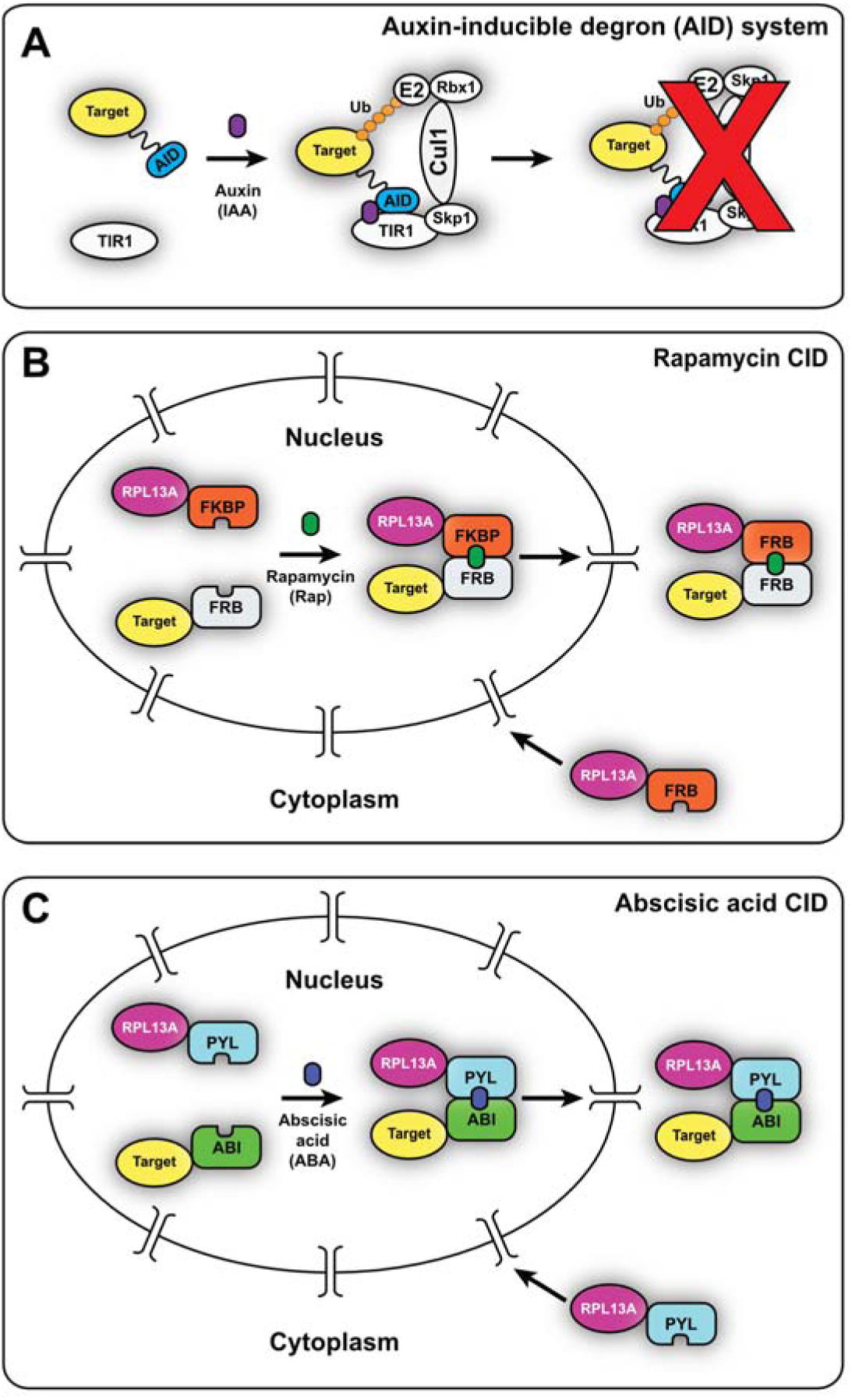
Schematic overview of chemical-induced dimerisation approaches for conditional depletion of nuclear proteins. (A) The auxin-inducible degron (AID) system requires the AID epitope to be fused to the protein of interest. Upon auxin addition, the TIR1 auxin receptor binds the AID degron tag. Together, TIR1 and the AID-tagged target form the substrate-binding platform of the E3 ubiquitin ligase SCF complex (i.e., Skp1, Cul1, and Rpx1 in the diagram). In turn, this assembly recruits an E2 ligase, which polyubiquitylates the AID degron tag and ultimately promotes the proteasomal degradation of the target protein. (B) In the rapamycin-dependent anchor-away (AA) technique, the Rpl13A-FKBP anchor protein is synthesised in the cytoplasm and rapidly imported into the nucleus shortly afterwards. There, it is incorporated into pre-ribosomal particles (not depicted in the diagram). In the presence of rapamycin, the anchor forms a tight complex with FRB-fused target proteins and, when associated with maturing pre-60S complexes, drags the target protein into the cytoplasm, where it remains sequestered on translating ribosomes. (C) Similarly to its rapamycin-mediated counterpart, the ABA-AA system fuses the PYL domain to Rpl13A, which also acts as the anchor in this procedure. In this case, abscisic acid (ABA) serves as a chemical inducer triggering dimerisation of the PYL-tagged Rpl13A to the target protein, which must have been previously modified to integrate an ABI epitope.

The AA technique relies on ligand-induced export of the target protein to the cytoplasm, where it remains sequestered on ribosomes or membrane-bound factors ^14^. This procedure leverages the ability of the human 12 kDa FK506 binding protein (FKBP12) and the 11 kDa, FKBP12-rapamycin-binding (FRB) domain of human mTOR to form a stable heterodimer in the presence of rapamycin ^16^ (Fig. 1B). In the original version of the protocol, the target protein is marked with the FRB domain, and the FKBP12 domain is genetically fused to the ribosomal protein Rpl13A ^17^, allowing the latter to act as a highly abundant cytoplasmic anchor (Fig.1B). Following its translation, Rpl13A is imported into the nucleus, where it is incorporated into 60S pre-ribosomal particles (Fig.1B). Bound to almost fully mature 60S particles, Rpl13A is exported back to the cytoplasm, where it then contributes to protein biosynthesis (Fig.1B). Therefore, in the presence of rapamycin, the target-FRB protein is dragged out of the nucleus and sequestered to translating ribosomes in the cytoplasm (Fig.1B).

Despite proving tremendously powerful, each of these CID methods have significant disadvantages. The original AID system was hindered by the cytotoxicity of auxin. The hormone can also undergo photodecomposition into harmful derivatives ^18^ that, if accumulated, cause growth defects. Additionally, the method displayed basal or ‘leaky’ degradation of the target protein as TIR1 weakly associates with the tagged target even in the absence of auxin, leading to unintended proteasomal degradation ^19^. These problems have been largely alleviated by substituting auxin with indole-free analogues^18^ that are more stable and less toxic, or by the AID2 methodology, which uses bump-and-hole engineering to prevent basal proteolysis and vastly reduce the ligand concentration required to induce target depletion ^20^. Regardless, the expression of the TIR1 protein still needs to be carefully tuned, as too high levels can induce uncontrolled breakdown of the target protein even in the absence of auxin ^15,20,21^.

The primary liability of the rapamycin-dependent AA system is its reliance on rapamycin. In addition to being expensive, this drug can bind to the yeast homolog of FKBP12 (Fpr1) to form an inhibitory complex that blocks the kinase activity of Tor1, which regulates cell growth, metabolism, and nutrient responses as a key component of the TOR signalling. To circumvent this problem, the AA technique must be applied to rapamycin-resistant strains encoding a mutant version of *TOR1* (*tor1-1*) and lacking *FPR1* (Δ*fpr1*). Being the most abundant FK506-binding protein in yeast ^22^, Fpr1 must also be removed to ensure that it will not compete with the FRB-tagged targets for the available FKBP12-fused anchors ^23^. As well as being technically challenging to implement, this required genetic background (*tor1-1* and *fpr1Δ*) diminishes the cell’s capacity to activate the heat-shock response ^24^. We are studying the role of RNA-binding proteins Nrd1-Nab3-Sen1 in regulating rapid adaptive responses in yeast upon changes in nutrient availability ^25,26^. Accordingly, the rapamycin AA strategy is not ideally suited to investigating these proteins that orchestrate transcriptional reprogramming of its targets under these stress conditions ^26–30^.

To address some of the problems associated with these CID systems, we have developed a modified AA system that exploits the plant phytohormone abscisic acid (ABA) to trigger dimer formation between the target protein and its cytoplasmic anchor ^31^ (Fig. 1C). In plants, ABA docks into the PYL domain of the RCAR, a receptor of the ABA signalling pathway, enabling the latter to associate with the ABI domain of the ABI1 protein phosphatase (reviewed in ^32^). This ABA-dependent CID event has been harnessed to build a synthetic spindle checkpoint in fission yeast ^33^. Inspired by this study, we recognised the potential of ABA in an AA system: it is inexpensive, non-toxic, and, since it engages in relatively weak interactions with PYL, its effect can be readily reversed by simply washing out the hormone ^31,33,34^ (Fig.1C).

Here, we demonstrate that ABA-dependent CID can be effectively employed as an AA system to deplete nuclear proteins in *Saccharomyces cerevisiae*. We show that ABA-AA rapidly and reversibly sequesters the abundant nuclear protein Nab3 in the cytoplasm and can deplete other highly abundant proteins, including the proteasome component Rpn11. Compared to the rapamycin-dependent variant, the ABA-AA requires higher doses of the ABA ligand to deplete the target protein and elicit the accompanying phenotypic effects. Nevertheless, we demonstrate that, contrary to other ligands used, ABA is innocuous, as it does not significantly impair cellular growth or dramatically alter gene expression in exponentially growing cells, even at very high concentrations.

We conclude that the ABA-AA system is highly complementary to existing tools. Unlike rapamycin-based AA, it has no toxicity issues and is fully reversible, making it ideal for studying dynamic processes. Critically, requiring only two fusion proteins generated using the provided integrative vectors, the ABA-AA system is readily applicable to diverse genetic backgrounds, environmental conditions, and other eukaryotic organisms.

## Results and Discussion

### ABA-AA recapitulates Nab3 depletion without toxicity in *S. cerevisiae*

Others and we have previously employed the rapamycin AA system to successfully deplete a variety of nuclear RNA surveillance factors, demonstrating the effectiveness of this approach ^26,35–38^. However, as it relies on the *tor1-1* and *fpr1Δ* mutations, this procedure may plausibly affect the TOR signalling pathway, which is intimately linked to nutrient sensing ^39,40^. We reasoned that introducing disturbances in that metabolic route was not ideal when examining the function of proteins implicated in adaptation to nutrient starvation. One such factor is Nab3, an essential RNA-binding protein involved in transcription termination that orchestrates transcriptional reprogramming during glucose or nucleotide shortage. Hence, we wanted to devise an AA system that would allow us to deplete Nab3 from the nucleus without having to interfere with the activity of TOR kinases.

Firstly, we began testing an optimised yeast auxin-induced degron (AID) system ^15^ (Fig. 1A). Briefly, this procedure required us to fuse the AID domain to the C-terminus of Nab3 and transform the BY4741 reference strain with a plasmid encoding a β-estradiol-inducible version of *TIR1* (pZTRL). To check whether this procedure could trigger the growth defects associated with loss of Nab3, we exposed the resulting Nab3-AID + pZTRL strain to different growing conditions: (i) the absence of any ligands (Figure 2A; ‘Untreated’); (ii) a medium containing a mix of auxin and β-estradiol, wherein *TIR1* would be expressed and able to induce degradation of the AID-labelled target; (iii) plates containing β-estradiol, which would merely induce *TIR1* expression; and (iv) the presence of auxin, which, by itself, would build up and moderately impair cell growth (Fig. 2A). To confirm these expectations, we monitored the growth of a strain in which the splicing factor Prp22 had been previously depleted with the same AID system (Fig. 2A) ^15^. Conversely, the BY4741 reference strain was used as a negative control to assess the wild-type yeast’s inherent tolerance to auxin and heterologous *TIR1* expression (Fig. 2A).

**Figure 2.**
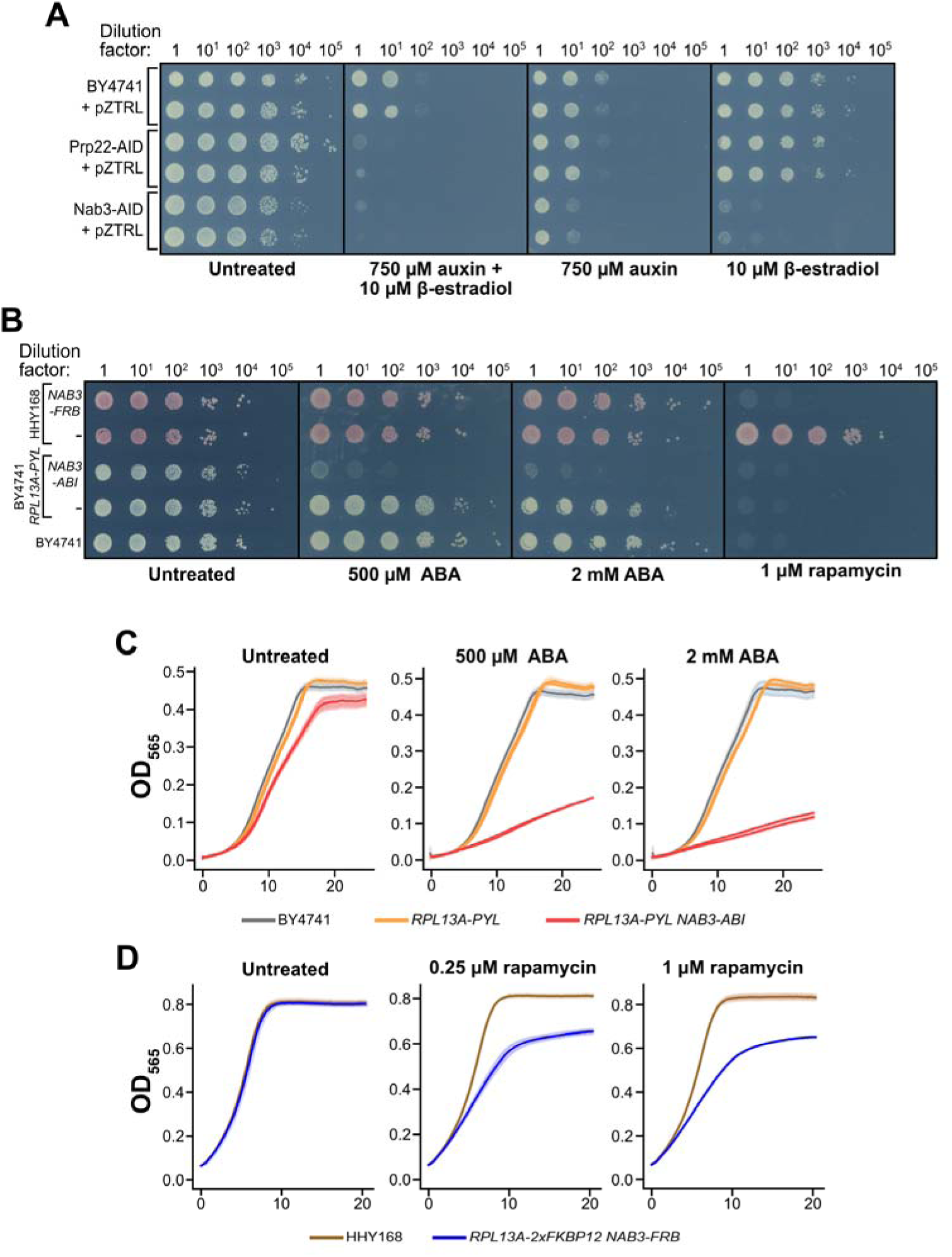
ABA is a non-toxic inducer that drives controlled degradation of Nab3 as efficiently as reported AID or rapamycin-dependent systems. (A) Representative images of the spot test analyses testing the efficiency of the AID system in selectively degrading Nab3 in the *NAB3*-*AID*. *NAB3*-*AID* and *PRP22*-*AID* contained a pZTRL plasmid encoding the TIR1 auxin receptor under a β-estradiol-inducible promoter. *PRP22*-*AID* was included as a positive control, whereas the BY4741 reference strain served as a negative control. Serial dilutions of cells were spotted onto plates containing minimal medium supplemented with glucose and the indicated doses of auxin and/or β-estradiol. (B) Representative images of the spot test analyses testing the efficiency of the ABA and rapamycin-dependent AA systems in selectively degrading Nab3 in the *NAB3*-*ABI* and *NAB3*-*FRB* strains, respectively. The BY4741 reference strain was included to as a negative control. Serial dilutions of cells were spotted onto plates containing minimal medium supplemented with glucose and the indicated dose of ABA or rapamycin. (**C**-**D**) Time-resolved optical density measurements depicting the mean (line) and the standard deviation (shading) of three technical repeats. Two independent clones were examined for each ABA-dependent AA strain.

As anticipated, the AID system efficiently degraded Prp22-AID in an auxin- and *TIR1*-dependent manner (Fig. 2A). However, β-estradiol-induced *TIR1* expression alone caused substantial growth defects in the Nab3-AID strain, indicating auxin-independent degradation. (Fig. 2A). This finding reemphasises the importance of carefully fine-tuning *TIR1* induction and production ^15,21^. Of note, tuning *TIR1* expression remains a caveat even in improved systems like AID2, where the auxin ligand-related drawbacks have almost been eliminated ^20^. Addition of auxin alone caused significant growth defects, and this was slightly exacerbated in Nab3-AID (Fig. 2A). We propose that, combined with the administration of auxin, leaky expression of *TIR1* from the pZTRL plasmid resulted in uncontrolled proteolysis of Nab3-AID.

We aimed to develop a conditional nuclear depletion system that was simpler and had less toxicity issues. We evaluated the potential of an abscisic acid (ABA) based CID system. ABA is innocuous to cultured mammalian cells ^31^, inexpensive, chemically stable, and has been applied as a dimerisation inducer in other yeasts ^33,34^. Thus, to test whether ABA was indeed harmless to *S. cerevisiae*, we monitored the growth of the BY4741 reference strain in media containing high concentrations of the hormone (i.e., 500 µM and 2 mM, 2 and 8 times more than the tested concentration in fission yeast ^33^; Figs. 2B-C). Our assays show that none of the ABA concentrations tested noticeably impaired cell growth (Figs. 2B-C), suggesting that at even at concentrations in the millimolar range, ABA does not significantly impact *S. cerevisiae* growth.

Next, we assembled the genetic parts needed for a functional ABA-dependent AA system. We constructed a strain expressing a hemagglutinin (HA)-tagged PYL domain (*PYL-3XHA*; 26 kDa) fused to the ribosomal protein Rpl13A (*RPL13A-PYL*). Next, in the *RPL13A-PYL* strain, we then C-terminally fused the FLAG-tagged ABI domain (*ABI-3xFLAG*; 37.4 kDa) to Nab3 (*NAB3-ABI*). We also generated a strain with reversed domain architecture (*RPL13A-ABI* and *NAB3-PYL*). However, Rpl13A-ABI was non-functional as an anchor. We speculate this could be due to the ABA-ABI-PYL interaction geometry or different epitope sizes (25 kDa vs 37.4 kDa).

Because the ABI and PYL domains comprised relatively large epitopes, we tested whether the target and anchor proteins remained fully functional. Even though *RPL13A* is a non-essential gene, tagging it with the 26 kDa PYL-3HA domain could affect the activity of Rpl13A and, in turn, inhibit translation to some exent. On the other hand, even mild deficiencies of Nab3 termination can severely disrupt cellular physiology ^25^. After performing growth assays in agar plates and liquid culture, we determined that introducing the *PYL-3xHA* tag downstream of Rpl13A did not significantly alter the growth of *RPL13A-PYL* with respect to the BY4741 reference (Figs. 2B-C). However, it should be noted that the microplate reader growth measurements revealed that integrating the ABI domain to the C-terminus of Nab3 modestly reduced the growth of *NAB3-ABI* (*RPL13A-PYL-3xHA NAB3-ABI-3xFLAG*) compared to the BY4741 reference strain (Fig. 2C; ‘Untreated’). Although such defect is not observed in the *NAB3-FRB* (*RPL13A-2xFKBP12 NAB3-FRB*) strain, some C-terminal fusions are known to reduce Nab3 function slightly ^30^. Nevertheless, the fact that this growth defect was not immediately apparent from spot-test analyses underscores the mildness of the defect (Fig. 2B; ‘Untreated’). Therefore, we conclude that the genomic integrations required for our ABA-dependent AA system largely preserved the function of the target and anchor proteins.

We then assessed whether fusing *ABI* and *PYL* to the target (i.e., Nab3) and the anchor (i.e., Rpl13A) allowed us to reproduce the growth defects associated with Nab3 loss in the presence of rapamycin ^30,38,41^. We found that 500 µM ABA caused growth defects in *NAB3-ABI* comparable to those of *NAB3-FRB* treated with 1 µM rapamycin (Figs. 2B-D), indicating near-complete Nab3 depletion. Interestingly, 500 µM of ABA constitutes twice the concentration used in previous studies in fission yeast and murine cells ^31,33^. However, Nab3 is highly abundant and, accordingly, may require higher ABA doses for depletion compared to less abundant proteins (Table 1). To test whether higher ABA concentrations could deplete more Nab3 from the nucleus, we tracked *NAB3-ABI* growth in 2 mM ABA. Since this barely exacerbated the growth defect (Figs. 2B-C), we concluded that 500 µM ABA achieves near-maximal Nab3-ABI depletion.

**Table 1.**
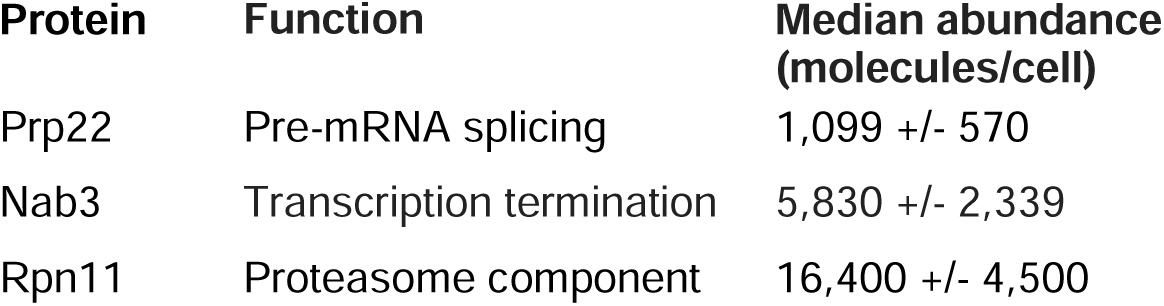
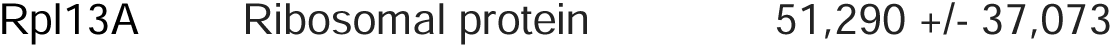
Median molecules per cell according to the unified protein abundance dataset generated by Ho *et al*.^59^.

The ABA-induced growth defect persisted for 24 hours (Fig. 2C), confirming ABA stability in liquid medium at 30°C. Together, these results demonstrate that ABA is an innocuous, inexpensive, and stable dimerisation agent that recapitulates rapamycin-induced Nab3 depletion phenotypes.

### ABA only marginally impacts gene expression in *S. cerevisiae*

Although 2 mM ABA (8 times higher than prevous studies; ^31,33^) did not noticeably affect cell growth, we tested whether it alters gene expression by comparing transcriptomes of BY4741 grown in minimal medium ± ABA (500 µM or 2 mM). BY4741 cultures were grown in minimal medium with 1% methanol (control), 500 µM ABA, or 2 mM ABA. After 5 hours of exponential growth, cells were harvested for RNA sequencing and data processed using the pyCRAC pipeline^42^, and a statistical comparison between treated samples and the methanol control was performed using DESeq2 ^43^ (Fig. 3 and Supplementary Table S1).

**Figure 3.**
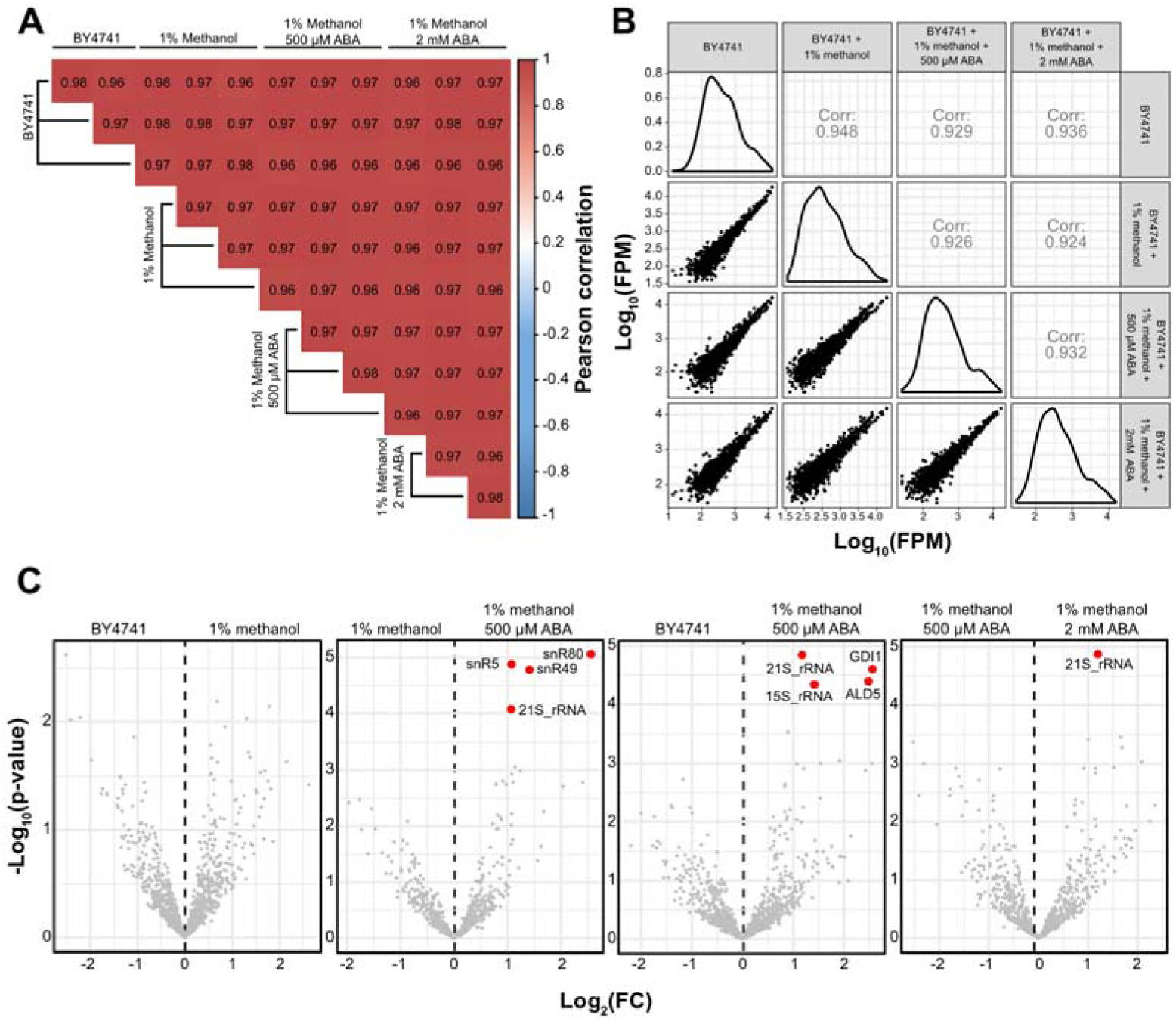
ABA treatment modestly alters gene expression profiles in *Saccharomyces cerevisiae*. (A) Matrix listing the Pearson correlation coefficients between the three independent experiments conducted under each condition (i.e., untreated ‘BY4741’, ‘1% methanol’, ‘500 μM ABA’, and ‘2 mM ABA’). (B) Comparison of BY4741 gene expression levels under the tested contexts. Panels comprising the main diagonal display density plots of log-transformed fragments per million (FPM) values for each experimental regime. Charts below the density plots show pairwise scatter plots of gene expression values across treatments. Frames above the main diagonal report the Pearson correlation coefficients between the transcriptomic profiles of each sample group. (C) Volcano plots summarising the outcome of the differential expression analysis conducted on the RNA-sequencing datasets obtained for each biological replicate under the tested conditions. Highlighted proteins (in red) were deemed differentially expressed: log_2_(fold-change) of more than 1 and a false discovery rate (FDR) of <= 0.05.

Read counts for genomic features across the three replicates were highly correlated (Fig. 3A), suggesting that the independent experiments generated reproducible results. Likewise, comparison of mean read coverage for genomic features in treated and untreated samples showed that treating cells with ABA or methanol did not have major impacts on gene expression (Figs. 3B-C). However, ABA treatment did significantly increase expression of several mitochondrial ribosomal RNAs (i.e., 21S rRNA and 15S rRNA), and two genes encoding a mitochondrial aldehyde dehydrogenase (*ALD5*) and a GDP dissociation factor (*GDI1*). In addition, ABA treatment increased the levels of three small nucleolar RNAs (snoRNAs; snR5, snR49, and snR80) (Fig. 3C).

The few differentially expressed genes showed no clear connection to known ABA effects. Notably, *ALD5* (encoding aldehyde dehydrogenase) was upregulated. While trace abscisic aldehyde (AB-ald, an Ald5 substrate and ABA biosynthesis intermediate) in our high-purity ABA (≥98%) could explain this, *ALD5* was similarly elevated in methanol-only cultures (fold-change:: 1.24; adjusted p-value = 0.45; Table S1). Since Ald5 oxidizes formaldehyde (produced from methanol by alcohol dehydrogenases) to formic acid, *ALD5* upregulation likely reflects methanol metabolism rather than ABA exposure.

Consistent with enhanced activity of the mitochondrial aldehyde dehydrogenase, upregulation of the mitochondrial ribosomal RNAs could equally stem from ABA treatment. In plants, ABA is known to alter mitochondrial redox balance, thereby affecting mitochondrial gene expression ^44^. Since mitochondrial redox signalling routes are largely conserved among species, it is plausible that such mitochondrial transcriptome changes are, at least partially, echoed in yeast. Ultimately, this could explain why ABA addition increases 21S and 15S rRNA levels, which are encoded within the mitochondrial chromosome. Gdi1 regulates the traffic of vesicles by controlling the dissociation of GDP from GTP-binding proteins ^45^. Interestingly, in rice, ABA has been shown to induce the synthesis of OsRhoGDI2, a stress-responsive GDI ^46^. However, the molecular basis underlying this phenomenon remains uncharacterised.

Thus, ABA concentrations up to 2 mM cause neither growth defects nor major transcriptional changes in budding yeast.

### The ABA-dependent AA system confers fully reversible depletion of Nab3

*NAB3-ABI* required 2000-fold more ligand than *NAB3-FRB* (500 µM ABA vs. 0.25 µM rapamycin; Figs. 2B-D), indicating that the rapamycin-FKBP12-FRB ternary complex is more stable than the ABA-PYL-ABI complex. This difference reflects the binding affinities underpinning each target-anchor interaction: the ABI-PYL interaction has a K_D_ of 52 µM in the presence of ABA ^47^, whereas the FKBP12-rapamycin-FRB ternary complex binds ∼4300-fold more tightly (K_D_ of ∼12 nM) ^16^. Despite requiring higher ligand concentrations, the weaker PYL-ABA-ABI interaction should enable full reversibility.

To confirm the concentration difference and compare depletion kinetics and reversibility, we performed time-resolved analysis of Nab3 nuclear depletion in both strains. We added ligand to induce depletion, then washed cells to remove the drug and assess reversibility. To quantify depletion efficiency and reversibility in this time-resolved assay, we exploited the NNS complex requirement for snR13 snoRNA termination and 3’ end processing ^48^ (Fig. 4A). When any NNS component is depleted, Pol II transcribes beyond the normal termination site, producing read-through transcripts that extend into the downstream TSR31 gene (Fig. 4A) and continues to synthesise species that extend to the downstream *TSR31* gene ^48^ (Fig. 4A). These *SNR13* readthrough products can be quantified by real-time quantitative PCR (RT-qPCR) reaction and serve as a proxy for Nab3 depletion efficiency. To determine Nab3 depletion kinetics, we measured 3’-extended snR13 levels for two hours following ligand addition (Figs. 4B-C). To assess reversibility, we washed out rapamycin or ABA and quantified snR13 read-through for an additional two hours (Figs. 4B-C).

**Figure 4.**
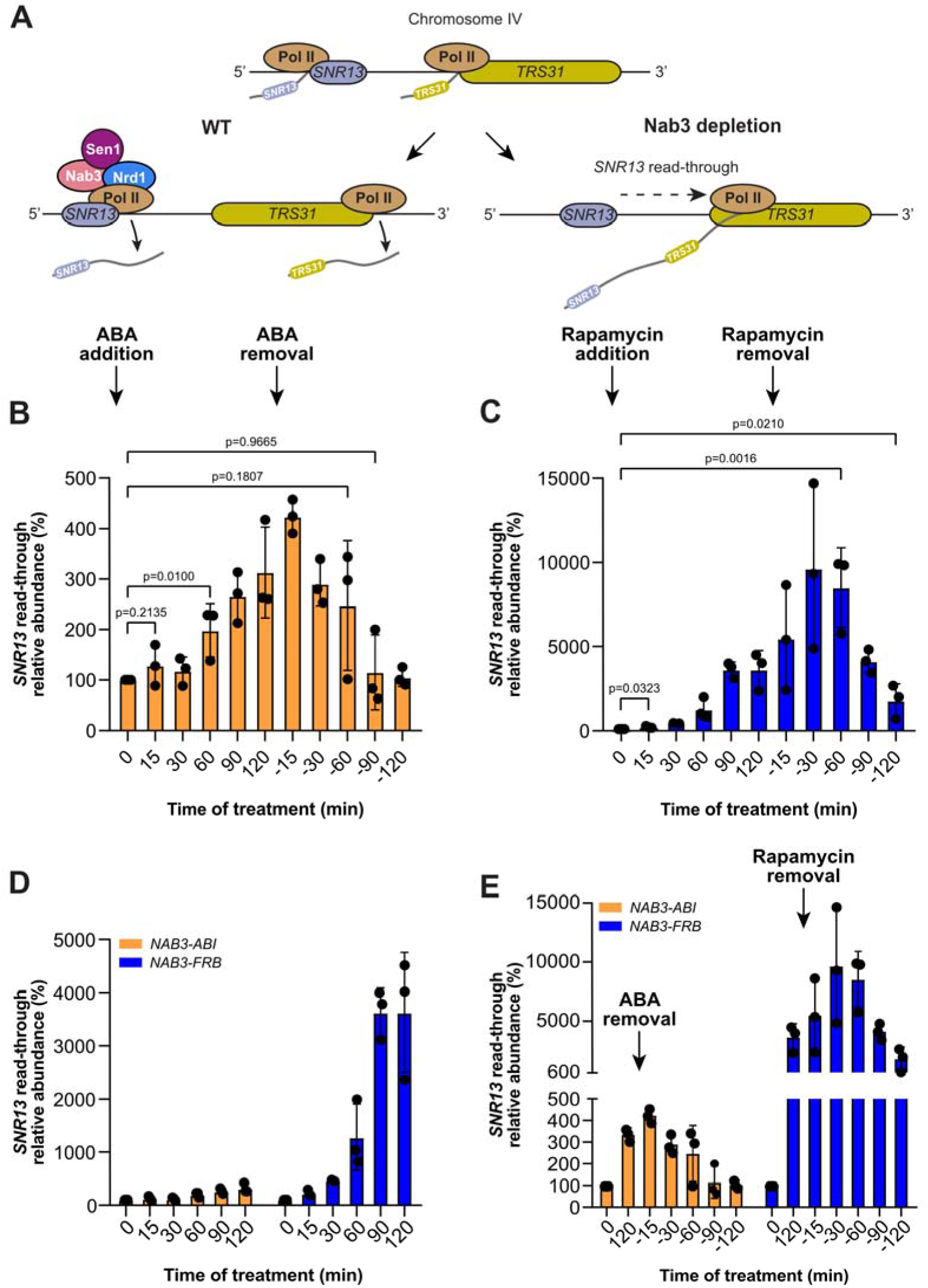
The ABA-dependent AA system enables fully reversible nuclear depletion of proteins. (**A**) Diagrammatic representation of *SNR13* snoRNA termination. *SNR13* (purple) and *TRS31* (green) loci lie on chromosome IV and are both transcribed by the RNA polymerase II (Pol II). The Nrd1-Nab3-Sen1 (NNS) complex binds to nascent *SNR13* snoRNAs and Pol II, promoting transcription termination and release of the snoRNA transcript. In this way, full assembly of the NNS complex onto their *SNR13* snoRNA targets prevents interference with downstream *TRS31* transcription. Loss of Nab3 results in transcriptional read-through of Pol II from *SNR13* into *TRS31*. This phenomenon yields extended RNA species comprising sequences from both *SNR13* and *TRS31*. (**B-C**) Relative abundances of 3’-extended *SNR13* transcripts were measured using RT-qPCR and normalised to the measurement recorded right before administering ABA or rapamycin (0 minutes). The inducer was added to cultures undergoing the mid-exponential growth phase, and RNA was isolated from samples gathered periodically. After 2 hours of treatment, the hormone and the drug were washed away from the pertinent cultures, and RNA was extracted from batches of cells collected at spaced intervals. Bar plot displays means and SDs from three independent experiments. P-values were calculated using a two-sided ratio paired t-test. (**D-E**) Bar graphs comparing the relative abundances of 3’-extended *SNR13* transcripts in the *NAB3*-*ABI* and *NAB3*-*FRB* strains upon addition (**D**) and removal (**E**) of ABA or rapamycin, respectively. As in B and C, the plot shows averages and SDs from triplicates. P-values were calculated using a two-sided ratio paired t-test.

Approximately one hour of ABA treatment was sufficient to detect significant accumulation of 3’-extended snR13 species in NAB3-ABI cells (Fig. 4B), whereas extended snR13 in *NAB3-FRB* was readily detected after 15 minutes of rapamycin treatment (Fig. 4C). Due to the higher affinity of the FKBP12-rapamycin-FRB ternary complex, rapamycin both depleted Nab3 faster and sequestered a greater fraction of the Nab3 pool than ABA (Fig. 4D). After two hours of treatment, extended snR13 levels in *NAB3-FRB* were seven-fold higher than in *NAB3-ABI* (Fig. 4D). However, despite more efficient Nab3 depletion, the rapamycin system showed minimal recovery: snR13 processing remained impaired two hours after drug removal (Figs. 4C and 4E). At this time-point, snR13 read-through levels in *NAB3-FRB* were six-fold higher than pre-treatment levels (Fig. 4E). Strikingly, snR13 read-through continued increasing until 90 minutes after rapamycin removal (Figs. 4C and 4E). In contrast, 3’-extended snR13 in *NAB3-ABI* began decreasing 15 minutes after ABA withdrawal and returned to pre-treatment levels within 1-1.5 hours (Fig. 4B).

These assays demonstrate that, despite being slower than the rapamycin-based system, ABA-AA is fully reversible, a valuable feature for studying nuclear protein dynamics. For instance, temporarily retaining a cytoplasmic pool of target protein provides an ideal starting point to study nuclear import kinetics. After inducing partial depletion with ABA, hormone removal allows the target to return to the nucleus and reimport can be tracked using live-cell fluorescence microscopy, such as fluorescence recovery after photobleaching (FRAP)^49^, which quantifies the rate and extent of molecular movement by monitoring how quickly fluorescence returns to a bleached region over time. It should also be possible to study transcription dynamics by temporarily titrating transcription factors and/or RNA polymerase subunits from the nucleus to the cytoplasm and following how transcription resumes after their controlled reintroduction. Another advantage of the ABA-AA system in this experimental setup is that, by gradually restoring the native distribution and concentration of the target protein, it (i) avoids saturating the nuclear import machinery and (ii) reflects the import-export balance more accurately than methods that fail to reinstate the original steady-state distribution (e.g., the poorly reversible rapamycin-mediated AA).

We previously reported that, when grown in medium containing a low concentration of a non-fermentable sugar, prolonged rapamycin-mediated sequestration of Nab3 in the cytoplasm causes significant cell size increases after several hours of rapamycin treatment ^25^. Consequently, we hypothesised that if ABA-AA were fully reversible, cell size should return to normal after ABA removal. To test this, we (i) monitored cellular volume in NAB3-ABI and *NAB3-FRB* cultures for 10 hours during ABA or rapamycin treatment, respectively; and (ii) reanalysed both cultures 10 hours after compound removal (the ’recovery phase’). We also evaluated cell size in two control strains: *RPL13A-PYL* (anchor only) and HHY168 (rapamycin-resistant parental strain) treated only with the corresponding solvents (i.e., 1% methanol and 1% ethanol, respectively). After 10 hours of recovery, *NAB3-ABI* and control strain *RPL13A-PYL* reached exponential phase, whereas *NAB3-FRB* remained growth-arrested (Fig. 5A), confirming rapamycin-based AA irreversibility. ABA-treated *RPL13A-PYL* grew identically to solvent controls, while rapamycin-treated HHY168 showed mild growth inhibition compared to ethanol-only controls, demonstrating rapamycin toxicity independent of target depletion. Thus, unlike rapamycin, ABA-AA is not toxic.

**Figure 5.**
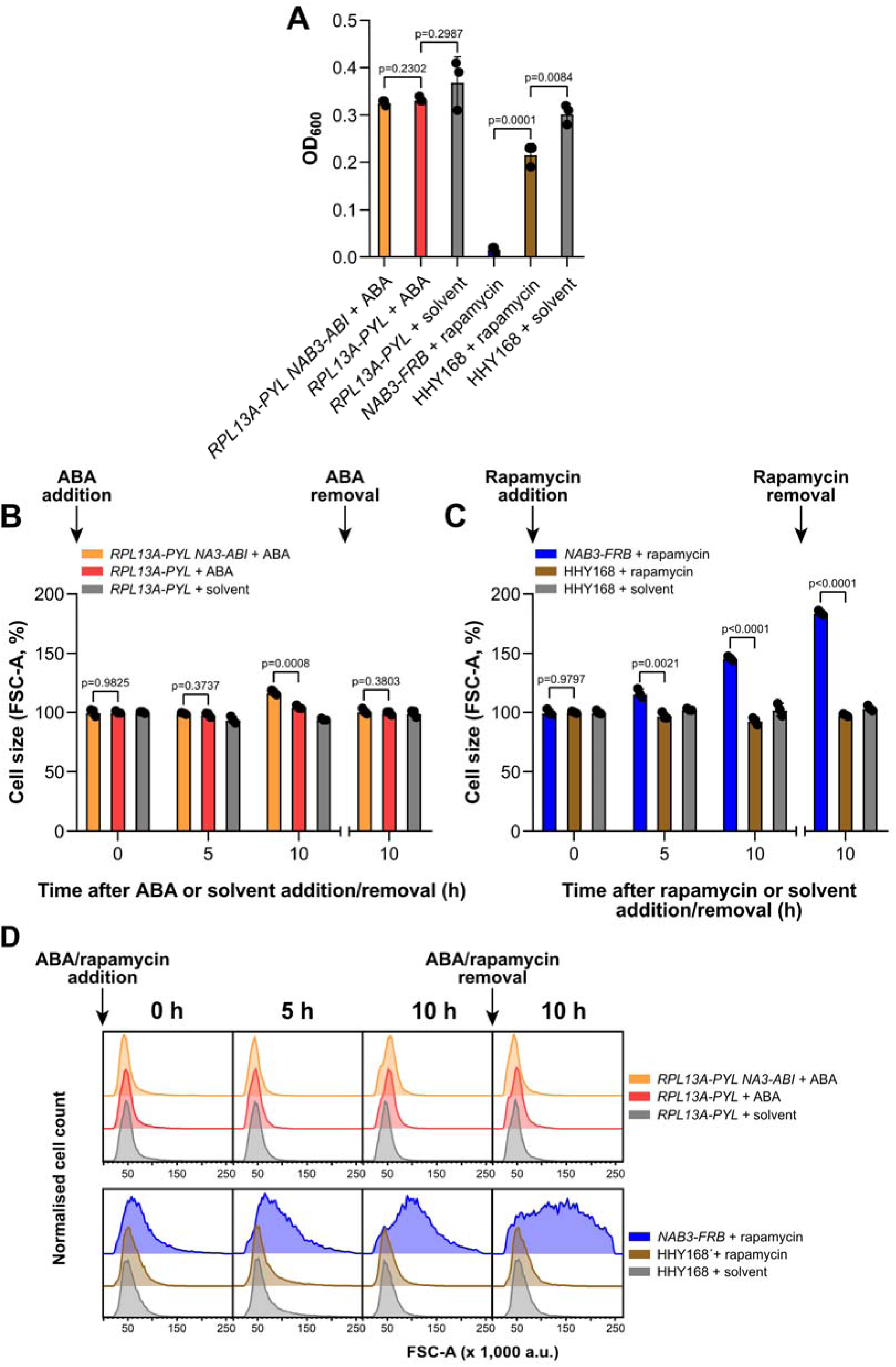
ABA-induced nuclear depletion is fully reversible. (**A**) Bar graph showing the recorded optical densities of the examined cultures of *NAB3*-*ABI* and *NAB3*-*FRB* and their parental strains 10 hours after washing away ABA or rapamycin from the medium. (**B-C**) Weighted means and SDs of the cell size (FSC-A values in flow cytometry) for three independent biological repeats of *NAB3-ABI*, *RPL13A-PYL*, *NAB3-FRB* and HHY168 were calculated for each strain and normalised to their volume before ABA, rapamycin or solvent addition. P-values were calculated using a two-sided unpaired Student’s t-test. (D) Representative flow cytometry traces of the cell size distribution of *NAB3*-*ABI* and *NAB3*-*FRB* and their parental strains treated with the indicated dimerisation agent or its solvent (Methanol or Ethanol) alone. The sizes were compared using their forward light scatter (FSC-A) values.

Flow cytometry showed *NAB3-FRB* volumes increased by 16% and 45% at 5 and 10 hours of rapamycin treatment. Unexpectedly, cells continued enlarging during recovery, reaching 80% above initial size (Figs. 5B-C), comparable to increases seen with conditional depletion of other NNS components ^25,29,50,51^. Since HHY168 showed no volume change throughout the experiment (Figs. 5B-C), we conclude that rapamycin itself does not cause cell enlargement in the rapamycin-resistant (*tor1-1 fpr1Δ*) background. Thus, the persistent size increase in *NAB3-FRB* confirms that the Nab3 rapamycin-based AA is not fully reversible.

*NAB3-ABI* cell size remained unchanged for the first 5 hours of ABA treatment but increased by ∼16% at 10 hours (Figs. 5B-C). Importantly, *NAB3-ABI* returned to normal cell size within 10 hours of ABA removal (Figs. 5B-C), confirming full reversibility of ABA-AA.

### The ABA-dependent AA system is applicable to abundant nuclear proteins

The effectiveness of any chemically induced dimerisation technique depends on the abundance of the target protein relative to that of the anchor. Rpl13A is present at around 50,000 molecules per cell, making it roughly 9 times more abundant than Nab3 (Fig. 6A and Table 1). Although Nab3 is among the most abundant proteins (80th percentile; Fig. 6A and Supplementary Table S2), we tested whether ABA-AA could deplete even more abundant proteins.

**Figure 6.**
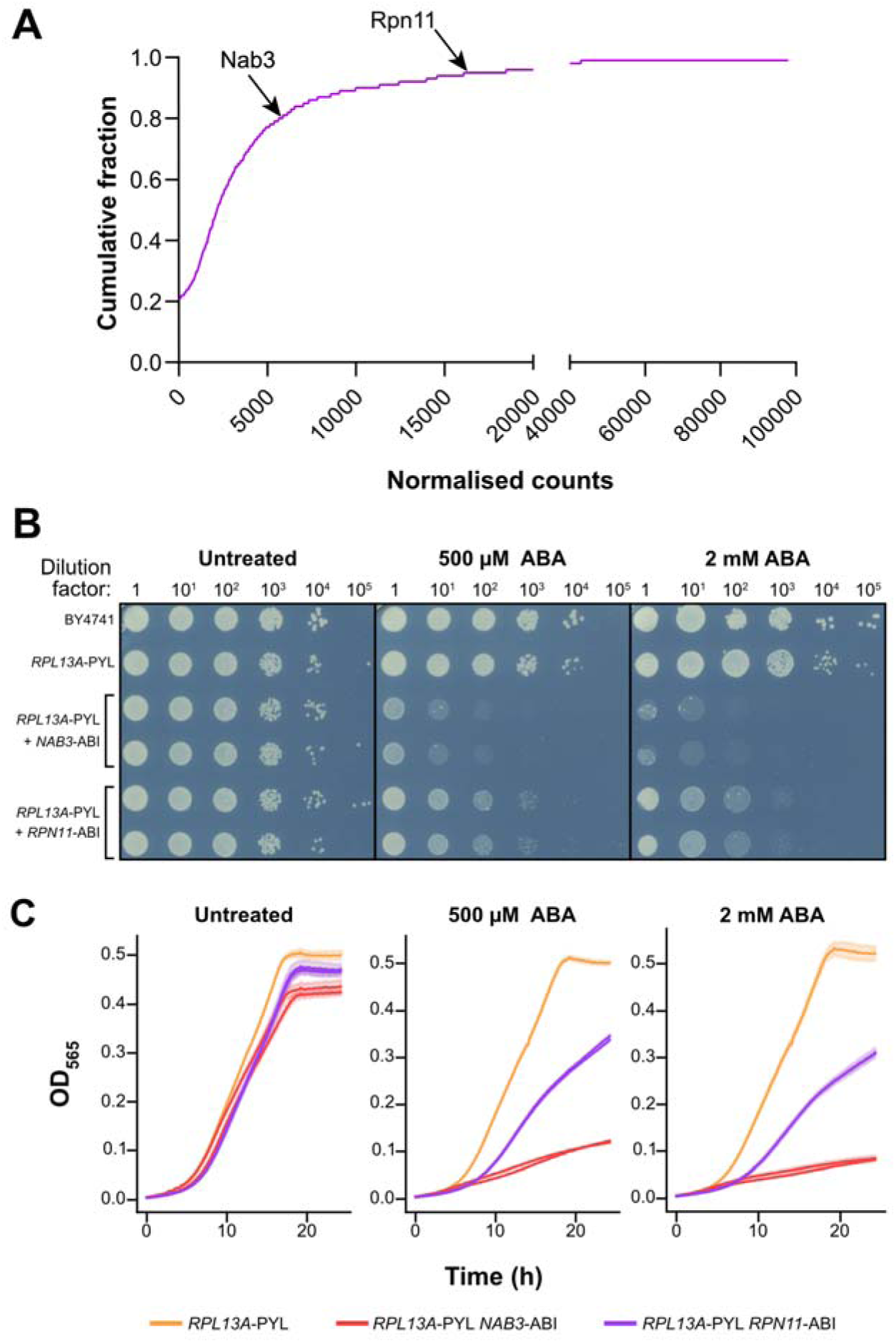
ABA-dependent AA can deplete even the most highly abundant nuclear proteins. (A) Nab3 and Rpn11 are highly abundant nuclear proteins. Cumulative distribution of molecules per cell for the nuclear proteome of *Saccharomyces cerevisiae* ^59,60^ (Table S2). Indicated in the plot are the abundances of Nab3 and Rpn11. (B) Representative images of the spot test analyses testing the efficiency of the ABA-dependent AA system in selectively degrading Rpn11 in the *RPN11*-*ABI* strain. *NAB3*-*ABI* served as a positive control for detecting defects in the presence of ABA. The BY4741 reference and the parental *RPL13A-PYL* strain were included as negative controls. Serial dilutions of cells were spotted onto plates containing minimal medium supplemented with glucose and the indicated dose of ABA or rapamycin. (**C**-**D**) Time-resolved optical density measurements depicting the mean (line) and the standard deviation (shading) of three technical repeats. Two independent clones were examined for the *RPN11*-*ABI* and *NAB3*-*ABI* strains.

We generated an *RPL13A-PYL* strain with ABI fused to the essential proteasome component Rpn11, which is ∼16,000 copies per cell (95th percentile; Fig. 6A, Table 1, and Supplementary Table S2). We then performed growth assays on agar plates and in liquid culture containing 500 µM or 2 mM ABA (Figs. 6B-C), with *NAB3-ABI* as a positive control and BY4741 as a negative control. *RPN11-ABI* grew normally without ABA, indicating that the tag does not impair function. However, 500 µM ABA depleted Rpn11 less efficiently than Nab3, requiring 2 mM ABA for substantial growth defects (Figs. 6B-C). This reinforces the importance of optimising ligand concentration for protein abundance. Since ABA induced substantial RPN11-ABI growth defects (Figs. 6B-C), ABA-AA can deplete even highly abundant proteins, providing a valuable alternative to rapamycin-based AA.

In summary, we have shown that ABA-AA efficiently depletes nuclear proteins in a fully reversible manner. Unlike rapamycin, ABA is non-toxic to wild-type yeast, does not dramatically alter gene expression, and is suitable for diverse genetic backgrounds. Requiring only two fusion proteins, ABA-AA provides a non-toxic and broadly applicable alternative for conditional protein depletion in budding yeast and other eukaryotes.

## MATERIALS AND METHODS

### Strains and growth conditions

*Saccharomyces cerevisiae* BY4741 ^52^ was the parental reference used to generate all the strains in this study. A list of all strains is provided in Table S3.

PCR-based modification was employed to fuse *ABI* domains downstream of NAB3 and *RPN11*; to integrate the PYL epitope distally to *RPL13A*; and to C-terminally tag *NAB3* and *PRP22* with AID, as previously described ^53,54^. The coding sequence of the *ABI* and *PYL* domains, as well as the AID tag, were amplified from the pFA6a-ABI-KanMX6, pFA6a-PYL-NatMX6, and pURA3-AID*-6FLAG vectors using the oligonucleotides listed in Supplementary Table S4 ^15,33^. Additionally, strains to be treated with the AID technology were transformed with the strain with the pZTRL plasmid, which codes for a β-estradiol-inducible version of *TIR1*. For clarity, a summary of the main features of each plasmid used for generating yeast strains is provided in Supplementary Table S5.

Yeast cells were grown in YPDA medium or synthetic complete (SC) minimal medium lacking tryptophan (SC -Trp) containing 2% (w/v) glucose as a carbon source. Agar plates of the same medium were prepared by supplementing the media with agar to a final concentration of 1% (w/v). While liquid cultures were incubated at 30°C with shaking at 190 rpm, cells grown in Petri dishes were incubated at 30°C in a static incubator for 48 hours, then kept at 4°C. Liquid and solid media was supplemented with 500 μM or 2 mM ABA, 0.25 or 1 μM rapamycin, 750 μM auxin and/or 10 μM β-estradiol.

All strains were pre-cultured in medium containing 2% (w/v) glucose for 24 hours prior to the start of the experiment. The cells used in all experiments were harvested during the exponential growth phase, except for those of the *NAB3-FRB* strain grown after rapamycin was removed from the medium (Figure 5D). For spot-test and microplate reader assays, cells were grown in 5 mL cultures. While RNA-sequencing was performed on extracts of cells collected from 25 mL cultures, real-time quantitative PCR and flow cytometry were conducted on samples harvested from 100 mL cultures. RNA was isolated from cells that were either pelleted by centrifugation (3202 rcf, 5 minutes), snap-frozen by submersion in liquid nitrogen, and stored at -80°C.

### Spot-test growth assays

Pre-cultures were diluted in 5 mL of SC -Trp to an OD_600_ of 0.1 and incubated for 3 to 4 hours at 30°C. Cells were then pelleted, washed with fresh medium, and resuspended in 1 to 3 mL of SC -Trp to reach an OD_600_ of 1. The resulting batches of cells were equally divided among the number of conditions (i.e., plates containing 500 μM ABA, 2 mM ABA, 1 μM rapamycin, 750 μM auxin, and/or 10 μM β-estradiol). Spot-test series were performed following 10-fold dilutions of the preceding culture. After inoculation with the relevant strains, plates were incubated at 30°C for 2 days.

### Microplate reader growth assays

Yeast cells from a culture in the mid-logarithmic growth phase were resuspended in fresh medium to a starting OD_565_ of 0.1. Two hundred µL was then transferred to three separate wells of a UV-sterilised 96-well black non-treated polystyrene microplate (Thermo Fisher Scientific) and its accompanying lid (Grenier). Triplicate controls comprising the same volume of sterile medium were also included for each examined condition. Absorbance was monitored in Infinite M200 (Tecan) microplate readers every 5-10 minutes at a constant temperature of 30°C while undergoing 535 seconds of orbital shaking at an amplitude of 6 mm between each recording time point.

Microplate reader data was analysed using the Omniplate package ^55^. The software corrected the OD measurements by subtracting the absorbance signal from the wells containing media only and compensated for the non-linear relationship between OD_565_ measurements and cell count at higher absorbance values. Corrected OD_565_ values were plotted using Python’s seaborn package.

### RNA extraction

RNA extracts from replicate samples generated from three different BY4741 colonies grown on different days. 5 mL SC -Trp cultures medium were incubated overnight before being used to inoculate 20 mL of SC -Trp, SC -Trp supplemented with 1% methanol, SC - Trp with 500 μM of ABA dissolved in 1% methanol, and SC -Trp containing 2 mM abscisic acid solubilised in 1% methanol. After 5 hours of growth, cells were harvested at an OD of approximately 0.5. RNA was purified following the established GTC-acid phenol extraction^56^

### RNA sequencing

RNA integrity was verified using the total RNA 6000 Nano kit on the 2100 Bioanalyzer (Amersham). Sequencing libraries from 3 biological replicates were generated using a modified version of the RNAtag-Seq protocol ^57^. One microgram of RNA from each sample was fragmented in FastAP phosphatase buffer for 6 minutes at 92°C in a final reaction volume of 20 µL. Subsequently, sheared RNA was dephosphorylated using 10 units of FastAP phosphatase and treated with 8 units of TURBO DNase™ (Thermo Fisher Scientific) in the presence of 40 units of rRNasin RNase inhibitor (Promega) for 30 minutes at 37°C in a 40 μL mix. Three prime adaptors (Table S4; ‘RNAtag_3adap’) were ligated to the dephosphorylated 3’ ends for 1 hour 30 min at 22°C in a 40 µL reaction volume with 1x T4 RNA ligase buffer, 9% DMSO, 10m ATP, 20% PEG8000 and 24 units rRNAsin (Promega). RNA was phenol-chloroform extracted and purified by ethanol precipitation.

Ribosomal RNA was subsequently removed using the yeast RiboZero Gold (Illumina) kit according to the manufacturer’s procedures. Free adapters were removed using Terminator Exonuclease according to the manufacturer’s procedure. The cDNA synthesis reaction was performed with Superscript IV following the manufacturer’s procedures at 55°C for one hour in a final volume of 14 µL. RNA was removed by adding 4 µL of 1N NaOH. A second ligation reaction was performed 3Tr3 DNA adapter (Supplementary Table S4) with CircLigase (Lucigen) according to the manufacturer’s procedures. Free adapters were subsequently removed using the Terminator™ 5-phosphate-dependent exonuclease (Thermo Fisher Scientific). The cDNA library was amplified by PCR using the P5 forward and barcoded reverse primers (2P_rev_1 and 2P_rev_2; Supplementary Table S4) for 24 cycles. PCR products were resolved on a 6% TBE gel (Invitrogen), and 180-400 bp fragments were gel excised. Final DNA concentration was measured using the Qubit 4 (Invitrogen) and the high-sensitivity DNA assay on the 2100 Bioanalyzer (Agilent).

### RNA-sequencing analysis

RNA sequencing was performed at NovoGene (Cambridge; 150bp PE). Raw FASTQ files were processed using the pyCRAC package (version 1.4.6 ^42^) using the CRAC_pipeline_PE pipeline ^26^. Briefly, adapter trimming was performed using Flexbar (version 3.4.0; ^58^) and reads were demultiplexed using pyBarcodeFilter.py. To remove PCR duplicates, reads collapsed using the pyFastqDuplicateRemover.py script. Subsequently, FASTA files with collapsed reads were aligned to the yeast genome (R64-1-1) using Novoalign (novoalign -d Saccharomyces_cerevisiae.R64-1-1.75.novoindex -f forward_reads.fasta reverse_reads.fasta -r Random; version 2.04; www.novocraft.com), and read counts were generated using pyReadCounters.py. Differential analysis of gene expression was performed using the DESeq2 R package ^43^.

### Real-time quantitative reverse transcription PCR

RNA extracts were purified using RQ1 RNase-Free DNase (Promega) and reverse-transcribed by SuperScript™ IV reverse transcriptase in combination with a random primer mix (New England Biolabs). Resulting samples were combined with Brilliant III ultra-fast SYBR® green qPCR master mix (Agilent Technologies), distributed within a 384-well plate (Roche), and amplified in a LightCycler® 480 qPCR instrument (Roche). *RDN5* and *PGK1* served as reference genes in all assays. Sequences for all RT-qPCR primers are listed in Supplementary Table S4.

### Cell size measurements using flow cytometry

1 mL from mid-logarithmic cultures was harvested, and isolated cells were fixed by the addition of 1 mL of cold 70% ethanol and incubated at -20°C overnight. On the following day, cells were permeabilised by two 10-minute incubations with 800 µL of 50 mM sodium citrate buffer (pH 7.2) at room temperature. RNace-It™ ribonuclease cocktail (i.e., a mixture of RNase A and RNase T1; Agilent Technologies) was then administered to each sample at a final concentration of 0.1 mg/mL before incubating them at 37°C overnight. Following RNA digestion, proteins were hydrolysed by adding 10 µL of 20 mg/mL proteinase K (Roche) and incubating the samples at 55°C for 2 hours. After sonication (5 seconds, 25 µm), samples could then either be stored for up to 1 month at 4°C or stained with 50 μg/mL propidium iodide (Sigma-Aldrich) for flow cytometry. For the latter instance, 200 µL of each set of cells were transferred into individual wells of a clear 96-well plate (Thermo Fisher Scientific) and analysed in a BD LSR-Fortessa (BD Biosciences) at the Flow Cytometry Facility for the School of Biological Sciences of the University of Edinburgh. A total of 10,000 events were acquired per sample.

### Flow cytometry analyses

Experimental files were exported in their native FCS format and analysed in FlowJo 10.8.1. Before proceeding with downstream analyses, we introduced a gate to select for single cells. For cell size experiments, the normalised distribution of FSC-A values was plotted in histograms. The weighted average and standard deviation were calculated across the three independent biological repeats assessed per cell type and condition. Strains were compared using unpaired t-tests run in GraphPad Prism 8.1, which was also used to generate the summary bar graphs.

## Data Availability

The sequencing data is available from the Gene Expression Omnibus (GEO accession GSE149746). The pyCRAC package and the pipeline used to analyse the RNA sequencing data are available from https://git.ecdf.ed.ac.uk/sgrannem/pycrac, from PyPi https://pypi.org/search/?q=pyCRAC and https://git.ecdf.ed.ac.uk/sgrannem/crac_pipelines

## Declaration of generative AI and AI-assisted technologies in the manuscript preparation process

During the preparation of this work the author(s) used Claude Sonnet in order to correct spelling and grammar mistakes. After using this tool/service, the author(s) reviewed and edited the content as needed and take(s) full responsibility for the content of the published article.

## Author Contributions

Conceptualisation, methodology, supervision and funding acquisition: S.G. Investigation: T.W., S.E-S., H.G. and I.F. Visualisation: S.E-S., T.W., H.G. and S.G. Writing - original draft: S.E-S. and S.G. Writing - review and editing: all authors.

## Acknowledgements

We are grateful to Isabella Maudlin, Jean Beggs, Manu Shakla, Priya Amin, and Kevin Hardwick for providing reagents and plasmids, and for fruitful discussions. We thank the members of the Swain and Granneman laboratories for critically reading the manuscript.

Flow cytometry data were generated within the Flow Cytometry and Cell Sorting Facility in the Ashworth Building at the University of Edinburgh. The facility was supported by funding from the Wellcome Trust and the University of Edinburgh.

This work was supported by a Medical Research Council Programme grant (MR/Y013131/1) to S.G., a SynBio-Med Wellcome Trust institutional fund to S.G., a Wellcome Trust PhD training fellowship (224084/Z/21/Z) to S.E-S., and an Erasmus+ traineeship to T.W.

